## Supplementary files 1a and 1b for "Sensory experience controls dendritic structure and behavior by distinct pathways involving degenerins"

**This file includes:**

Supplementary file 1a
Supplementary file 1b

Supplementary References

**Supplementary file 1a.** List of strains and transgenes used in this work

| Strain | Genotype | Details |
|---|---|---|
| N2 | Wild-type | (1) |
| JPS282 | *asic-1(ok415) I*; *vxEx282[WRM0621dC07 + unc-122p::GFP]* | CGC |
| VC244 | *gtl-1(ok375) IV* | CGC |
| SS104 | *glp-4*(*bn2*) *I* | CGC (2) |
| CB1338 | *mec-3(e1338) IV* | CGC (3) |
| MT1085 | *unc-8(n491) IV* | CGC (4) |
| VC2633 | *degt-1(ok3307) V* | CGC (5) |
| DR466 | *him-5(e1490) V* | CGC |
| NC279 | *del-1(ok150) X* | CGC (5) |
| CB1611 | *mec-4(e1611) X* | CGC (6) |
| ZB2551 | *mec-10(tm1552) X* | CGC (5) |
| JPS478 | *asic-1(ok415) I*; *mec-10(tm1552) X*; *vxEx478[sto-5p::asic-1(+) unc-122p::GFP]* | CGC |
| AQ3272 | *ljEx637[PF49H12.4::DEGT-1::mCherry Punc-122::GFP]* | Provided by W. Schafer (5) |
| AQ3273 | *ljEx638[PF49H12.4::MEC-10::mCherry Punc-122::GFP]* | Provided by W. Schafer (5) |
| ZX819 | *lite-1(ce314) X*; *zxIs12*[*pF49H12.4::ChR2::mCherry pF49H12.4::GFP*] | Provided by A. Gottschalk (7) |
| BP709 | *hmnIs133(ser-2Prom3::Kaede)* | Provided by M. Heiman (13) and used by T. Gattegno (8) |
| BP925 | *mec-4(e1611) X; hmnIs133(ser-2Prom3::Kaede)* | Cross (9) |
| BP1021 | *him-5(e1490) V*; *hmnIs133(ser-2Prom3::Kaede)* | Cross: BP709 X DR466 |
| BP1022 | *mec-10(tm1552) X*; *hmnIs133(ser-2Prom3::Kaede)*; *him-5(e1490) V* | Cross: BP1021 X JPS478 |
| BP1023 | *asic-1(ok415) I*; *him-5(e1490) V; hmnIs133(ser-2Prom3::Kaede)* | Cross: BP1021 X JPS478 |
| BP1024 | *asic-1(ok415) I; mec-10(tm1552) X*; *him-5(e1490) V; hmnIs133(ser-2Prom3::Kaede)* | Cross: BP1021 X JPS478 |
| BP1025 | *asic-1(ok415) I; degt-1(ok3307) V*; *mec-10(tm1552) X* | Cross: BP1024 X VC2633 |
| BP1026 | *degt-1(ok3307) V; mec-10(tm1552) X* | Cross: BP1024 X VC2633 |
| BP1027 | *degt-1(ok3307)* V; *hmnIs133(ser-2Prom3::Kaede)* | Cross: BP1021 X VC2633 |
| BP1028 | *asic-1(ok415) I; degt-1(ok3307) V; hmnIs133(ser-2Prom3::Kaede)* | Cross: BP1025 X BP1027 |
| BP1029 | *mec-10(tm1552); degt-1(ok3307) V; hmnIs133(ser-2Prom3::Kaede)* | Cross: BP1025 X BP1027 |
| BP1030 | *asic-1(ok415) I; mec-10(tm1552) X; degt-1*(*ok3307*) *V*; *hmnIs133(ser-2Prom3::Kaede)* | Cross: BP1025 X BP1027 |
| BP1031 | *degt-1(ok3307) V*; *ljEx638[PF49H12.4::mec-10::mCherry Punc-122::GFP*] | Cross: VC2633 X AQ3273 |
| BP1033 | *mec-10(tm1552) X*; *ljEx637[PF49H12.4::degt-1::mCherry Punc-122::GFP]* | Cross: BP1022 X AQ3272 |
| BP1034 | *mec-10(tm1552) X*; *hmnIs133(ser-2Prom3::Kaede); him-5(e1490) V; hyEx321[ser-2Prom3::mec-10genomic]* | pWRS825 plasmid provided by W. Schafer (5) was injected into BP1022 |
| EB1982 | *dzIs53[pF49H12.4::mCherry] II* | Provided by Y. Salzberg (10) |
| TV17924 | *wyIs50007[ser2prom3*::GCaMP6 *egl-17::mCherry] X* | Provided by K. Shen (11,12) |

**Supplementary file 1b**. List of primers used in this work

| Gene | Sequence of the primer |
|---|---|
| *asic-1(ok415)* I | *Forward-1: 5' aactggtgtggccacttcaactttc 3';*<br>Forward-2: 5' aaggtttcagatgatcgcgtagtcaag 3';<br>Reverse: 5' catttctcttcttccgtcagcgc 3' |
| *mec-10(tm1552)* X | Forward-1: 5' acacggctccttcttgagttccga 3';<br>Forward-2: 5' attcggtttcctcctcttcttccaatgc 3' ;<br>Reverse: 5' cgtttttttcagcgccctttcctgca 3' |
| *degt-1(ok3307)* V | Forward-1: 5' cgagtagctgattatcaaaaagtcctcga 3';<br>Forward-2: 5' cggatattccagcattggcgaa 3';<br>Reverse: 5' ttccccgttgatcttctatgtattaca 3' |

**Supplementary References**


1. S. Brenner, The genetics of *Caenorhabditis elegans*. *Genetics* **77**, 71-94 (1974).
2. M. J. Beanan, S. Strome, Characterization of a germ-line proliferation mutation in *C. elegans*. *Development* **116**, 755-766 (1992).
3. J. C. Way, M. Chalfie, *mec-3*, a homeobox-containing gene that specifies differentiation of the touch receptor neurons in *C. elegans*. *Cell* **54**, 5-16 (1988).
4. N. Tavernarakis, W. Shreffler, S. Wang, M. Driscoll, unc-8, a DEG/ENaC family member, encodes a subunit of a candidate mechanically gated channel that modulates *C. elegans* locomotion. *Neuron* **18**, 107-119 (1997).
5. M. Chatzigeorgiou *et al.*, Specific roles for DEG/ENaC and TRP channels in touch and thermosensation in *C. elegans* nociceptors. *Nat Neurosci* **13**, 861-868 (2010).
6. M. Driscoll, M. Chalfie, The *mec-4* gene is a member of a family of *Caenorhabditis elegans* genes that can mutate to induce neuronal degeneration. *Nature* **349**, 588 (1991).
7. S. J. Husson *et al.*, Optogenetic analysis of a nociceptor neuron and network reveals ion channels acting downstream of primary sensors. *Curr Biol* **22**, 743-752 (2012).
8. M. Oren-Suissa, T. Gattegno, V. Kravtsov, B. Podbilewicz, Extrinsic Repair of Injured Dendrites as a Paradigm for Regeneration by Fusion in *Caenorhabditis elegans*. *Genetics* **206**, 215-230 (2017).
9. V. Kravtsov, M. Oren-Suissa, B. Podbilewicz, The fusogen AFF-1 can rejuvenate the regenerative potential of adult dendritic trees by self-fusion. *Development* **144**, 2364-2374 (2017).
10. N.J. Ramirez-Suarez *et al.,* Axon-dependent patterning and maintenance of somatosensory dendritic arbors. *Dev Cell* **48**, 229-244 (2019).
11. Y. Cho, D.A. Porto, H. Hwang, L.J. Grundy, W.R. Schafer, H. Lu, Automated and controlled mechanical stimulation and functional imaging: In vivo in C. elegans. *Lab on a chip*, **15**, 2609 – 2618 (2017).
12. Y. Cho, D.N. Oakland, S.A. Lee, W.R. Schafer, H. Lu, On-chip functional neuroimaging with mechanical stimulation in Caenorhabditis elegans larvae for studying development and neural circuits. *Lab on a chip* **18**, 601–609 (2018).
13. Z.C Yip,. Heiman MG (2016) Duplication of a Single Neuron in C. elegans Reveals a Pathway for Dendrite Tiling by Mutual Repulsion. Cell Reports 15, 1–9
